# Temporal resolution shapes inferred higher-order interactions and information compression in neural population activity: A data-driven approach using minimally complex models

**DOI:** 10.1101/2025.03.23.644813

**Authors:** Fariba Jangjoo

**Affiliations:** Kavli Institute for Systems Neuroscience, Faculty of Medicine and Health Sciences, Norwegian University of Science and Technology, Trondheim, Norway

## Abstract

Studying higher order interactions in complex interacting systems based on limited data is inherently challenging. An additional, often overlooked factor is the temporal resolution at which data is presented for analysis. This study examines how temporal resolution influences the emergence of higher-order statistical dependencies and information representation in population level neuronal activity. Using a minimally biased data driven statistical inference framework, Minimally Complex Models, we studied grid-cell populations in the medial entorhinal cortex across a broad range of temporal resolutions. Results demonstrate that at intermediate temporal resolutions (100 milliseconds), the inferred model captures significant higher order dependencies alongside efficient information representations, whereas coarse resolutions lead to poor informative representation. These findings establish temporal resolution as a decisive factor in the inference of higher order neural structure and highlight that resolution-aware modeling is crucial for accurate analysis and characterization of collective dynamics in complex neuronal systems.

## Introduction

Understanding the role of higher-order interactions is crucial in order to study how complex interacting systems such as neural systems encode and process information. Traditionally, studies have focused on pairwise interactions, which capture the correlations between two elements. However, making prior assumptions about the underlying interaction structure is a biased approach when dealing with complex systems. By assuming pairwise interactions, deeper structure within interacting systems where higher-order interactions simultaneous correlations among multiple elements, is overlooked. Higher-order interactions offer a more informative understanding about neural dynamics and functional connectivity [1, 2]. In addition, consequences such as a compact form of encoding of information and processing are not effectively explained by simpler models such as pairwise models [3]. Studies indicate that pairwise models inadequately represent complex systems, as these fail to capture the rich dynamics of higher-order interactions in large neural populations [4, 5]. For instance, Ince et al. (2009) showed that interactions beyond pairwise, are essential in order to capture the complexity of the neural responses in the somatosensory cortex, where pairwise models alone are not able to properly approximate the information carried by the population of neurons [2]. Similarly, Zylberberg and Shea-Brown (2015) observed that nonlinearities in input could lead to higher-order correlations, improving information transmission in neural populations [6]. Other studies confirmed that despite weak pairwise correlations, higher-order correlations reveal deeper structure within data which emerge as highly coordinated network states [7]. In addition, studies showed the necessity of the higher-order interactions in understanding widespread distributions of neural activity [8, 9].

The ability to retain information from neural codes is also dependent on the temporal resolution of analysis or the temporal resolution that the data is recorded in. This can highly impact the interpretations of neural dynamics. Fine temporal resolution is able to capture detailed and rapid neural dynamics, whereas coarse resolutions may smooth out dynamical patterns, leading to information loss [10–13]. For example, Panzeri et al. (1999) illustrated how stimulus-independent correlations can contribute to information synergy in short time windows, emphasizing the need to choose appropriate temporal scales for accurate modeling [14]. However, many of the studies ignore the influence of time resolution on the significance of these types of interactions, often assuming that higher-order interactions are inherently tied to specific states without adjusting for dynamic temporal factors [15]. This assumption can lead to an incomplete picture that does not incorporate the fact that information content in neural spike trains is highly affected by the temporal resolutions that the system is studied in [16–18].This important factor impacts functional dependencies among elements and, as a consequence, the efficiency of encoding of information.

Higher-order interactions have also been linked to various co-variates such as brain states. Studies show that higher-order interactions are more prominent during specific conditions, such as heightened synchrony or attention. For example, research has demonstrated that neural networks exhibit stronger higher-order correlations during synchronized states like slow-wave sleep, compared to more asynchronous activity seen in awake or resting states [19, 20], while other papers show that higher-order interactions in the brain become more pronounced during the resting state [21, 22]. Higher-order interactions have also been shown to improve the neural population code’s ability to perform tasks like pattern recognition and decision-making [23]. In the visual cortex, research demonstrates that during active perception or directed attention to a stimulus, higher-order correlations can enhance the brain’s ability to discriminate between visual features [11, 24]. This enhanced discrimination may arise from the brain’s use of higher-order interactions to create distinct representations of stimuli, thereby increasing the efficiency of encoding and processing information. Similarly, in sensory systems, higher-order correlations become more pronounced during states of rich sensory input or heightened perceptual engagement. For example, Shlens et al. observed that in the primate retina, multi-neuron firing patterns exhibit stronger higher-order interactions under complex visual stimuli compared to simpler or less dynamic conditions [25]. This trend is also observed in the visual cortex, where Reynolds and Heeger (2004) reported enhanced discrimination between visual features due to higher-order interactions during active perception [24]. This suggests that higher-order interactions may be critical for encoding and processing information during complex sensory experiences.

However, inferring higher-order interactions is a challenging task especially when working with great number of elements. Traditionally, approaches such as maximum entropy models that attempt to account for higher-order interactions struggled with computational demands as the number of interacting elements grows [3, 26–28]. Moreover, these models rely on predefined assumptions about the significance of interaction orders, which can restrict compatibility when it comes to explaining complex systems based on limited data [5, 29]. Also, studies highlight the importance of incorporating higher-order interactions to accurately model neural population activity in practical applications, such as brain-machine interfaces, visual processing or in general when it comes to modeling biological systems [30–32].

In this paper, with the help of a data-driven approach that is able to explain the dependency of higher-order interaction on temporal resolution in neural systems, we aim to understand the underlying model that is able to reflect the configuration by which the data was generated. Unlike conventional approaches, the model is able to reveal which interaction orders are most relevant for a given dataset without predefined structural assumptions. By examining extensive data considering various neuronal geometries which result from spacing among cells, external and internal states which can be consequence of internal state of the animal (e.g., sleep) and configuration of the experiment, this approach reveals how higher-order correlations emerge differently across temporal resolutions underscoring that informational content and interaction significance are inherently linked to temporal framing. Moreover, our approach identifies an optimal time resolution where a minimal number of components encode the maximum amount of information, with higher-order interactions playing a dominant role. Our approach provides a structured framework in which understanding the generative model purely based on data is possible. Moreover, our study emphasizes that temporal resolution is not merely a pre-processing detail, rather it fundamentally influences the interaction structure inferred from neural population data.

## Materials and methods

### 0.1 Inference framework for binary data from complex interacting systems

In order to study the interactions defined by a binary dataset recorded from a complex interacting system, we employ the framework of Minimally complex models (MCMs). This framework is designed for binary data where the response of each system variable is simplified into two states. Given a sequence of binary dataset, the temporal resolution is defined as the bin size that summaries the response of the system variable. For simplicity, if the system variable is responsive somewhere in a particular bin with specific size (The method of determining the activity of the variables is defined in Binning procedure), the response of the variable is summarized as active. On the other hand, if the variable is non-responsive in that particular bin, the response of the variable is defined as silent.

For a system with *n* number of variables, a single observation of the system’s state can be defined as ***S*** = (*s*_1_, *s*_2_, …, *s*_*n*_), where each *s*_*i*_ represents the binary state of variable *i*. Therefore, a set of *N* independent observations of system state can be expressed as *S* = ***S***^(1)^, …, ***S***^(*N*)^. Given a system state ***S***, one relevant question to ask is about the underlying structure of the binary data of the system’s state, which defines the connectivity patterns among the variables in the system. Classically in the realm of statistical inference, the underlying model that the data is generated from is predefined. If we assume that the variables are connected two by two (pairwise interaction), the suggested underlying model could be pairwise Ising model. However, one can object over this assumption and suggest higher order interactions in the system as we do not have any precise information on the generative model. Therefore, in order to find the best structure for a given binary dataset, one must evaluate all the possible structures that can exist for a system with *n* number of variables. However, exhaustively evaluating all possible structures is an infeasible computational task. Therefore, according to the framework suggested by C. de Mulatier et. al. [33], we restrict our search to a particular family of models called minimally complex models (MCMs). According to this framework, the inferred structure has the minimal information theoretic complexity and at the same time maximizing the model’s evidence as measure of how well the model describes the data. If each particular structure is defined as a model, by comparing the evidence of models (posterior probability that the model generates the data, integrated over its parameter values), one can find the best structure as the model with the largest evidence.

The MCM framework assumes that a system configuration ***S*** is drawn from an exponential family distribution. In this framework, each variable *s*_*i*_ is binary (taking values of *±*1), and the probability of observing ***S*** is given by;

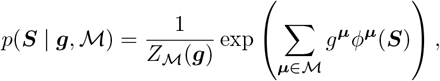

where ℳ represents the set of operators encoding interactions of arbitrary order, with each interaction defined by the function *ϕ*^***µ***^(***S***). Each operator ***µ*** indicates which variables participate in an interaction, represented as a binary vector or integer value of *n* bits. To be precise, operator ***µ*** can be represented as a binary vector with length *n*, or equivalently by a binary integer of *n* bits. For instance in a system with *n* number of spin variables, if a spin operator is product of three variables *s*_1_, *s*_4_ and *s*_5_, then the binary code (*µ*) of this particular spin operator is 000011001 or the integer representation 2^0^ + 2^3^ + 2^4^ = 25. Furthermore, *g*^***µ***^ denotes the interaction strength for operator ***µ***, and *Z*_ℳ_ (***g***) is the partition function ensuring normalization of *p*(***S***|***g***,ℳ).

In absence of any prior knowledge, one must evaluate all the possible models generated by *n* number of variables (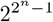number of models), which is an infeasible task. In order to tackle the computational challenge of searching among all the models, MCMs constrain the search such that the best models defined by orthogonal subsets called Independent Complete Components (ICCs).

Each ICC (denoted as ℳ_*a*_), is constructed from a set of *r*_*a*_ operators, which cannot be obtained by multiplying other existing operators in the set. This set is regarded as basis set. If the basis set consist of individual binary variables, then the basis of each ICC would include only those variables without any higher-order products (Cardinal *r* of basis set is equal to *n*). On the other hand, the basis set can be expanded to include combinations that cover maximal bias. In this case the cardinal *r* of basis set is less than. The later case results into 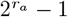 operators for each ICC, and consequently the number of operators for the entire model is defined as 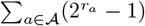. The combined basis set for the model ℳ is then can be obtained as the union of the bases ***b***_*a*_ for each ℳ_*a*_;

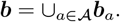

Each ICC ℳ_*a*_ is orthogonal to others, which means that the interactions within each component are independent of each others 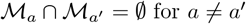 for *a* ≠ *a*^′^.

Based on Bayesian model selection, the best model is the model that maximizes the posterior probability *P* (*S* | ℳ). Using Bayes’ theorem *P* (ℳ | *S*) ∝ *P* (*S* | ℳ)*P*_0_(ℳ), one can attempt to calculate the evidence as

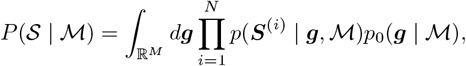

where *p*_0_(***g*** |ℳ) is the prior distribution over parameters.

If we define 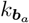 as the frequency of basis operator ***b***_*a*_ in the data set *S* with length *N*, then the model evidence for a given configuration ***S*** is calculated as [33]:

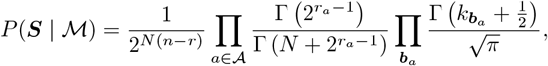

where Γ denotes the Gamma function. This formulation of the evidence enables comparison across MCMs, leading us to selection of the model with the highest evidence

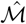 which is the best fit for the dataset.

Consequently, the maximum likelihood distribution will take the following simple format;

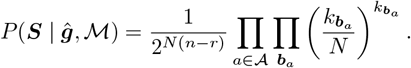

This formula allows simplify the exhaustive calculation of the log evidence, while avoiding over-fitting.

In summary, through MCM framework, the algorithm iteratively explores through the space of possible models, updating to models that lead to increase of the model evidence, until no alternative model yield to higher model evidence. Therefore, the final model 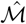 will provide an optimized description based on the dataset. Once the optimal MCM is found, further analysis can be done in order to explore attributes such as component sizes, interaction’s significance, etc.

### 0.2 Binning procedure

In order to evaluate the impact of the binning method on the results, two different binning methods are considered. Generally, binning involves dividing the entire duration of recorded data *T*, into *t* bins, where *t* is determined by the division of recorded length by the chosen bin size 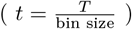. Thereafter, to summarize each bin’s data into a single representative value, we approached it by two different methods:

- **Presence of Spikes Method:** In this method, we assign a value of 1 to a bin if there is at least one instance of activity (i.e., the neuron fired) within that bin; otherwise, the bin is denoted as 0 to indicate inactivity.
- **Average Firing Rate Method:** In this approach, we compute the mean firing rate within each bin *R*_bin_, and compare it to the mean firing rate over the entire recording *R*_*T*_. If *R*_bin_ ≥ *R*_*T*_, the bin is marked as active (1), otherwise inactive (0).

### 0.3 Random realizations

In order to have better statistical evaluation, we need to generate random realizations of the recorded binary dataset. This procedure starts by specifying a fixed duration *T* for all realizations. Given this duration, we extract multiple sub-datasets (or “realizations”) of length *T* from the original dataset, which can be regarded as independent samples for further analysis. The analysis is done for various population sizes of cells *n*, where *n* is randomly chosen from the total cell population in the dataset. For each chosen cell population size *n*, we draw *N* = 200 independent realizations of the recorded dataset, each of length *T*, by randomly sampling subsets of cells from the entire dataset. Each realization is then binned according to Binning procedure, using a chosen bin size, which summarizes the activity within each bin. Thereafter, quantities of interest are calculated for these binned realizations at varying bin sizes.

In order to account for potential statistical biases due to limited data, we repeat the same analysis on a shuffled version of each realization, where before binning, the sequence of spikes for each cell is randomly permuted. Thereafter, the binning is done over the shuffled version of each realization. This shuffling is intended to remove any inherent temporal correlations while preserving the original firing rates. The results of the shuffled version of the realizations can be used to remove the bias due to limited data sampling. In order to do so, we evaluated the “difference” between the results of the shuffled and unshuffled versions. By examining the difference, we aim to isolate the effects of temporal correlations and other structured dependencies present in the original dataset, from random noise or baseline levels in the shuffled data. In figures, results from the original (unshuffled) data are denoted with the subscript *O* (Original), while results from the shuffled data are denoted with *S* (Shuffled).

### 0.4 Distribution of active cells across different bin sizes

After binning each random realization into a specific bin size, one can investigate the number of bins that have a specific fraction of active cells, or in other words, the probability of observing a fraction of active cells considering *N* data points. Clearly, in this procedure, when the bin size value is the finest (i.e. 1*ms*), each cell can only be active once. While considering the greater values of bin sizes leads to coverage of a longer time span, which might lead to coverage of multiple activities of one cell. In this analysis, we are not concerned about the number of spikes that each cell emits, rather **the presence of activity of each cell**. After counting the present cells in each bin, one can obtain the distribution of each fraction. In order to do so, the fraction of the bins with the occurrence of each fraction of active cells is counted. The distribution can be observed in a heatmap, where the y-axis shows the fraction of the active cells (*n*_*A*_), the x-axis shows the value of the bin size, and the heat color shows the probability of observing *n*_*A*_ having *N* data points defined by the bin size. This probability is evaluated over 200 different realizations of the data with a length of 10 minutes.

### 0.5 Independent complete components and log-evidence

After finding the best model (according to Inference framework for binary data from complex interacting systems) for each random realization at different bin sizes, we can perform analysis on the ICCs and the Log-evidence inferred from the inference framework. Having the ICCs and log-evidence of the inferred model, we investigate two main quantities; the normalized number of ICCs 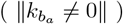 where this value is normalized by the number of cells *n*, and the log-evidence of the detected model 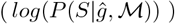 per data point *N* both normalized by *n* and unnormalized format. All these values are studied on average with their standard deviation, where these statistics are calculated considering all the random realizations. Bare in mind that the number of independent complete components is calculated by counting basis *b*_*a*_ which with 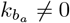. By normalizing the values to the number of cells *n*, we can obtain a better insight when comparing results of realizations with different values of *n*.

### 0.6 Orders of interaction

Another interesting attribute to evaluate is the order or equivalently the size of detected components and associated operators of the best model that describes the data. When observing the order of interactions, we need to perform two distinct procedures for components and operators. In order to evaluate the size of the components, the number of unique cells is considered, where this set of cells is obtained from the union of all the cells presented in all operators of the corresponding component. However, when dealing with operators, the size of all the operators of each ICC is evaluated.

Thereafter, in order to calculate the probabilities, we can investigate the number of components/operators with order *i* in which the components/operators contain *i* number of cells. One can ask how the fraction of components with order *i* changes by changing the value of the bin size. In order to see this behavior we tried to plot the fraction of components with order *i* within a model averaged over all the considered realizations for each bin size. However, the plots are not visually clear due to small values. In order to have visually clear plots, a cube root transformation was applied to the values. This transformation re-scaled the values by raising them to the power of 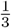, which reduces the impact of large outliers while retaining the relative differences between values. In heatmaps, the color bars show the respective original values, while the y-axis values in curve plots are presented in cube root transformation.

### 0.7 Dataset

In this study, the neural data that is used for analysis is the data recorded from grid cells in the medial entorhinal cortex [34]. The data acquisition by the means of tetrode arrays allowed for simultaneous recording of neural activity from multiple grid cells over a long period, which is essential for our type of analysis. We restricted our analysis to two rats renamed as R and Q, of which the data from rat R is recorded over two different days. In general, this recording includes sessions across various tasks, behavioral states, and cellular modules, which gives a richer understanding of dependencies on possible co-variates.

The recording is explained in more detail based on behavioral states which are indicated by task and state of the animal (e.g. during sleep), and modules which represent groups of grid cells that exhibit similar spatial scales in their firing patterns. Behavioral state or type of experiments are categorized as:

- **Open Field (OF):** A square-shaped environment where the rats could freely explore for randomly scattered food crumbs. Recordings were made to capture the hexagonal firing patterns of grid cells during active navigation.
- **Wagon Wheel (WW):** A circular track with radial arms, resembling a wagon wheel. This environment provides a structured spatial setting to study how grid cells adapt to a different spatial layout.
- **Rapid Eye Movement (REM) and Slow-Wave Sleep (SWS):** Recordings were performed to capture grid cell activity in two different sleep phases.

In addition, data for each rat is organized into modules, with each of these modules categorized based on the spatial scale of their grid patterns and their temporal firing characteristics. Furthermore, each module’s spatial configuration is characterized by a different level of grid spacing. Rat R is recorded across three modules which progressively, from module 1 to 3, increase the spacing between peaks in the hexagonal grid pattern. For rat Q, two modules are considered in which module one features closer spacing while the second one covers a larger spatial area. In this paper, the session’s recording is denoted as “Rat/Module/Experiment/The day of the recording”. In this setting, day one is encoded as I and day two as II. In the case that there is none, no day was indicated in the presented dataset. For instance, R1WWII is the recording from the first module of the rat R in WW experiment which is recorded in day 2. The detail of each recording is presented in table 1.

**Table 1.**
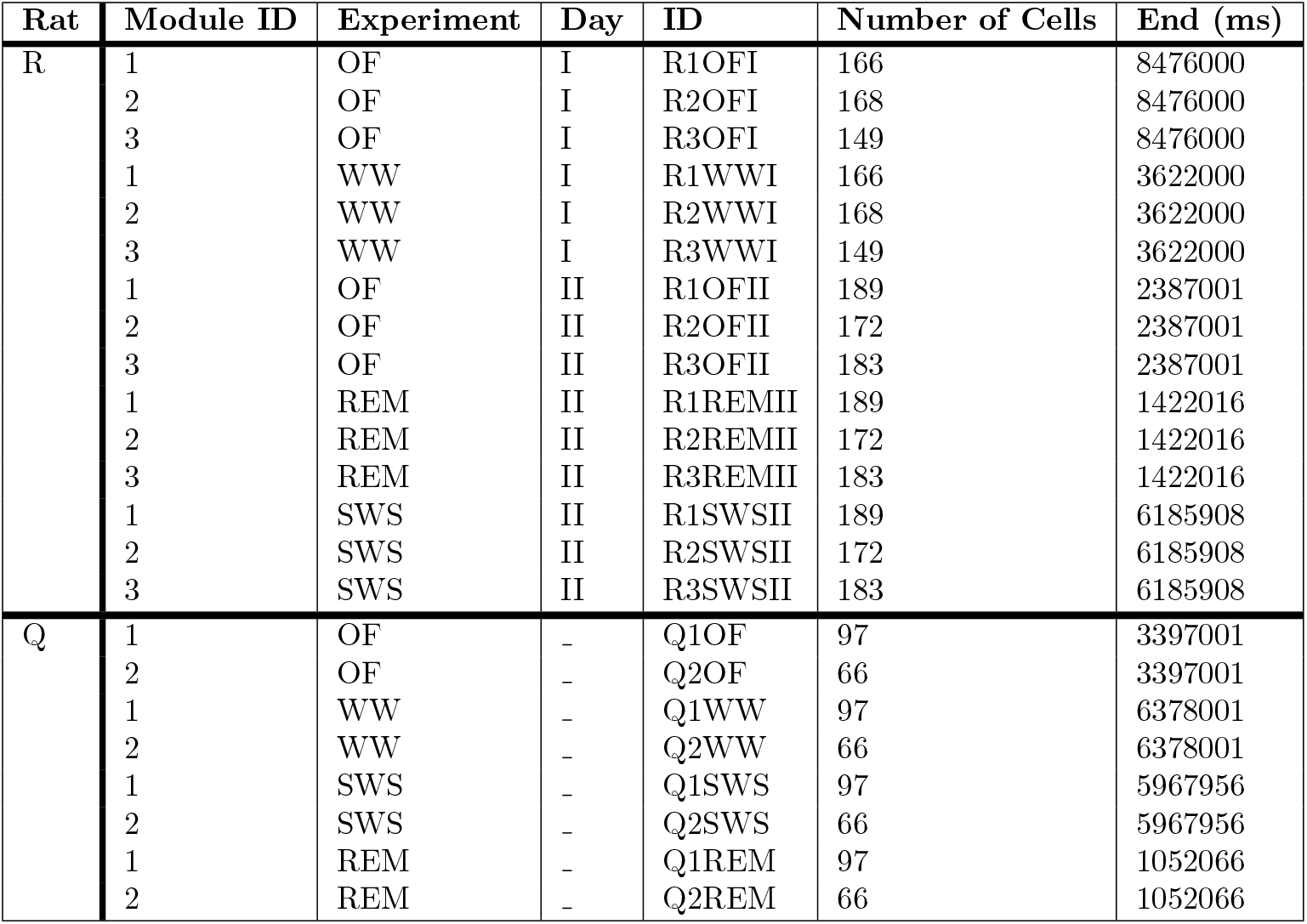
Information on the Grid cell’s recording of rat R and Q in 1 millisecond time resolution. This table summarizes the sessions, listing the rat name, Module ID, Experiment, the day of the recording, number of recorded cells and the total duration of each session at 1 millisecond temporal resolution.

## Results

In this paper, the results of the analysis on 23 distinct recordings of grid cells in the medial entorhinal cortex of two rats (R and Q) across different experimental states, days and modules, are extensively evaluated (Information of the recordings is presented in Table 1). From each recording, 200 random realizations with *N* number of data points (*N* = 10 minutes) and *n* number of cells are drawn independently. Thereafter, three types of analysis are performed; analysis on the realizations recording which is denoted by *O*, on shuffled realizations denoted by *S*, and the difference of these two types in order to understand the results by avoiding the statistical biases that come from limited data (Section 0.3). Thereafter, in order to understand the effect of the temporal resolution on our analysis, each individual realization’s data (Original or Shuffled), is evaluated for different values of bin sizes ranging from the finest resolution *b* = 2^0^ = 1 ms to the coarsest considered resolution *b* = 2^15^ = 32, 768 ms. The bin size sequence follows integer powers of 2 ({2^*i*^ | *i* ∈ {0, 1, 2, …, 15}}), with a few intermediate values (90, 181, and 724). In order to evaluate the effect of binning methods, the activity within bins is determined by two distinct methods; i.e. Presence of spikes and average firing rate methods (Section 0.2). Having the binned realizations, we attempt to infer the underlying model based on the minimally complex models MCMs (Section 0.1). All the analyses are evaluated for different populations of cells (*n* ∈ {10, 15, 20, 30}). In the following, you are presented with sample results belonging to one of the recordings (**R1REMII**) as there is no substantial difference in conclusions, especially when considering qualitative behaviors. Results for other recordings can be found in the public repository (https://osf.io/jn6m5/?view_only=33e6056f2f6a4ab396504344ccf9b57a). It is clear that the choice of temporal resolution can affect the observed activity patterns in a dataset. If we define the temporal resolution by the bin size that we attempt to bin a given binary dataset, then the smallest value of bin size allows at most one event per bin. While the greatest value of the bin size captures all the activity of a cell by one single bin. The smallest and the greatest values of the bin size for a recording with 10 minutes length are 1 millisecond and the length of the recording (10 minutes) respectively.

Another important factor to emphasize is the binning procedure as this factor can hugely affect the activity pattern of the cells within each bin. One can confirm that the method based on average firing rate implies stricter criteria in order to register a cell as active and this potentially leads to sparser binned data than the method based on the presence of spikes. In any of these cases, binning the data affects the temporal resolution that the data is analyzed, meaning that the decrease of the temporal resolution is equivalent to the increase of the bin size value. Consequently, the new spike patterns consist of unique information at each bin size value, which affects the inferred generative model.

In the following, we are going to understand important features of the generative models which give us a good understanding of the encoding of the information for different values of bin size and the dependence of the higher order correlation on the resolution that the data is looked at. By implementing the framework of MCMs, we obtained the independent complete components (ICCs) that can describe the given binary dataset and their associated operators through a heuristic search among all the possible models.

**ICCs:** One can consider the number of ICCs as the efficiency of the encoding of the information in the data as this parameter defines the number of needed parameters that can describe the data. When considering the number of ICCs inferred from the realizations of the dataset, one can see that the average number of the ICCs follows a curve with one clear global minima around ∼10^2^ milliseconds and a slight flatness or local minima which takes place around ∼10^4^ millisecond in the binning method based on average firing rate and slightly before ∼10^3^ in the binning method based on the presence of spikes (Fig 1A and Fig 1D). Evaluating the inferred models from both the original data and the corresponding shuffled data, shows that both have global minima, in which the global minima of the original data takes place at around the same value as the case where the difference of original data and corresponding shuffled data is considered (Fig 1C and Fig 1F), while for the shuffled case is around the same value as the local minima of the ICCs plot for the difference case (Fig 1B and Fig 1E). This can potentially tell us that these results are sensitive to the number of spikes and the inferred correlation among cells shows limited data bias; otherwise, one might expect a straight line pointing at absolute independence of cells for all values of bin size (in the case of absolute independency, the number of ICCs and the number of cells are equivalent, in which each ICC contains one cell). One can observe this effect in shuffled data for small values of the bin size, where the correlation is absent and therefore, the number of ICCs on average is equivalent to the number of cells, while the lack of data points in coarse temporal resolutions leads to false correlation and consequently fewer number of the ICCs. Another certain observation for all the cases of original, shuffled and difference is that the increase in the number of cells leads to fewer number of components up to the first global minima, which can show us the importance of including more number of cells as this can reveal richer structure within the data.

**Fig 1.**
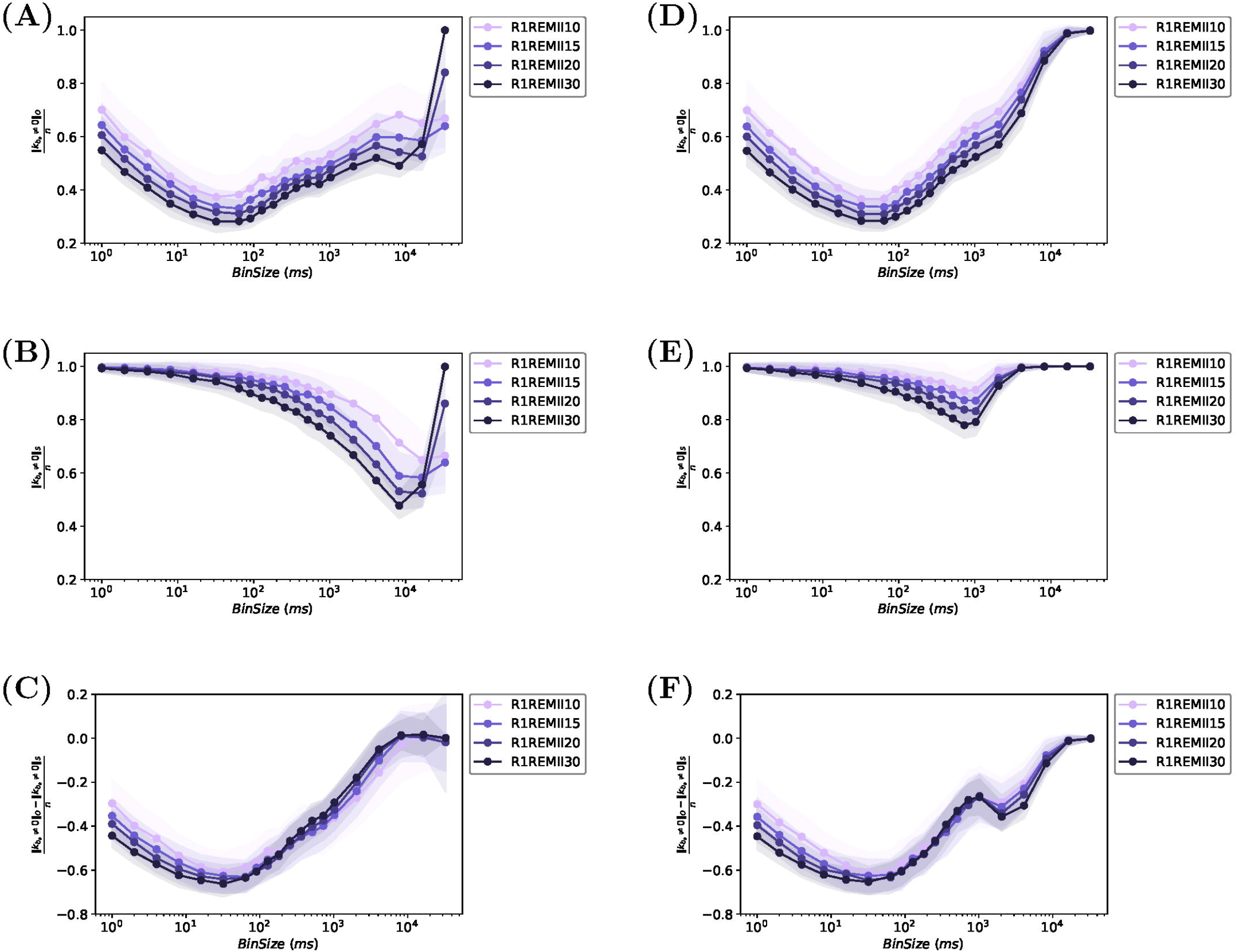
Normalized number of ICCs Vs. bin size for R1REMII dataset. The plots are shown for two different methods of binning where the plots in the left column are based on the average firing rate and the ones in the right column are based on the presence of spikes in bin. Plots for each binning procedure are shown for the original data set (panels A and D), shuffled data set (panels B and E) and the difference (panels C and F), where the difference of the number of the detected ICCs in the original dataset with the corresponding shuffled version is considered. All plots show a global minima regardless of the number of cells and the binning method. Moreover, the greater the value of the cell’s population leads to fewer numbers of detected ICCs. This pattern is clearer up to the global minima.

**Log-evidence:** By evaluating the log-evidence per data point of the detected models, one can confirm the same qualitative behavior (curves with minima) when considering the original and the corresponding shuffled cases, where the minima are centered around a value in [10^2^, 10^4^] (Fig 2A, Fig 2D, Fig 2B and Fig 2E).

**Fig 2.**
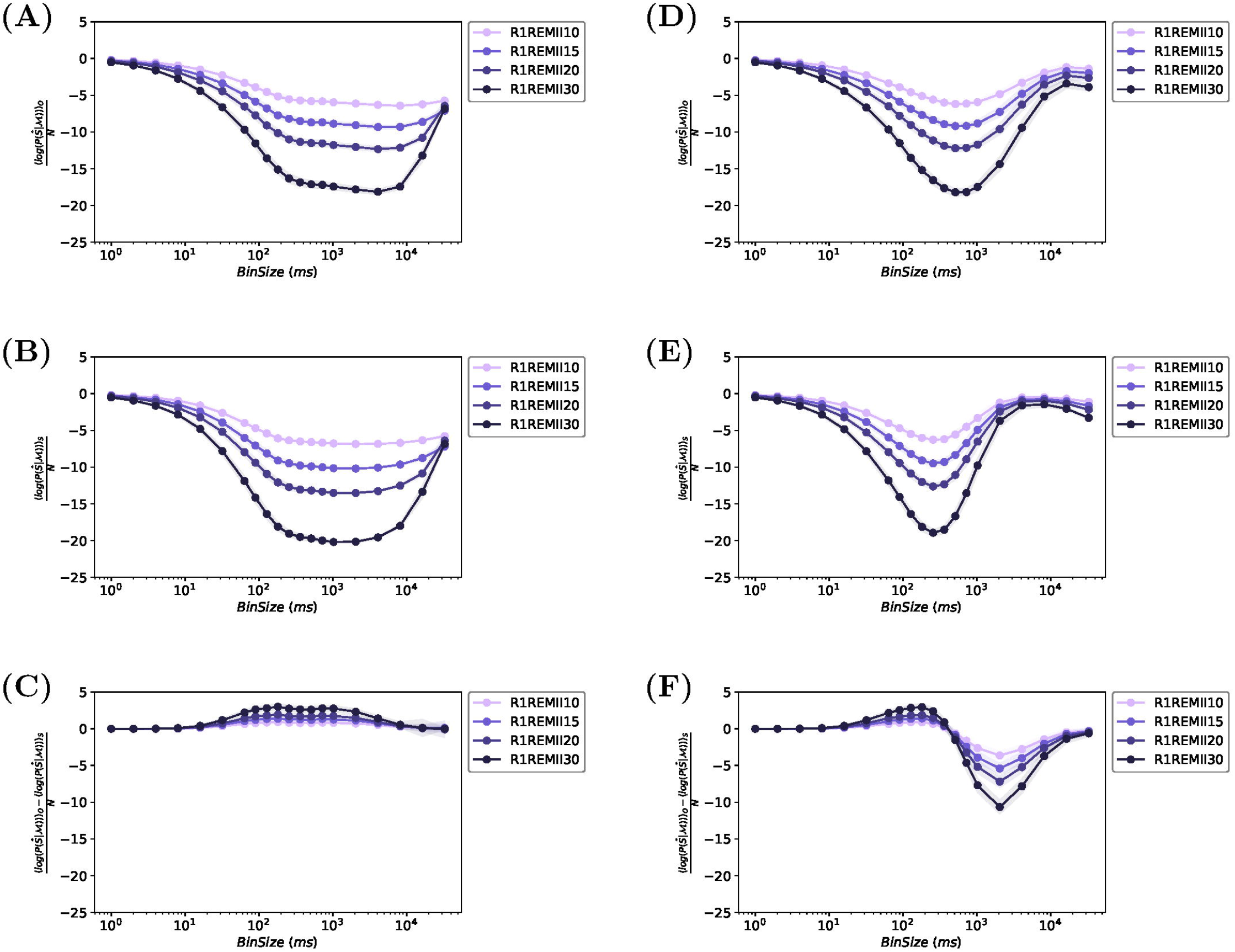
Log-evidence per data point Vs. bin size for R1REMII dataset. The plots are shown for two different methods of binning where the plots in the left column are based on the average firing rate and the ones in the right column are based on the presence of spikes in bin. Plots for each binning procedure are shown for the original data set (panels A and D), shuffled data set (panels B and E) and the difference (panels C and F), where the difference of the log-evidence per data point for the detected ICCs in the original dataset with the corresponding shuffled version is considered.

However, one can get a clearer picture by evaluating the difference of the original and corresponding shuffled case. These figures can potentially tell us for which range of bin size the log-evidence per data point is higher when avoiding the bias due to limited data. The difference plot shows a high value of log-evidence per data point around 10^2^ milliseconds regardless of the binning method (Fig 2C and Fig 2F). However, one can observe that the binning method based on the presence of spikes shows the poorest models according to log-evidence per data points by introducing false correlations due to a lack of data points. This false correlation increases as the number of cells increases, which leads to poorer models (Fig 2C and Fig 2F). However, this effect appears in greater bin sizes in the binning method based on average firing rate, which can be seen by the gradual decrease of log-evidence after ∼10^2^ milliseconds. Later on, we present the probability of active cells at different temporal resolutions, which can shed light on why this is observed.

In addition, if one attempts to normalize the log-evidence per data points curves by the number of cell 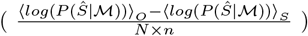, all the plots overlap (Fig 3). This can provide us a universal measure when it comes to the comparison of the models for different co-variates where the results are independent of the system size. Notice that 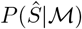 is equivalent to 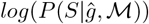 in our notations. The notation is the same with the logic that for a specific chunk of dataset 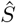, a set of parameters 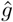 are detected.

**Fig 3.**
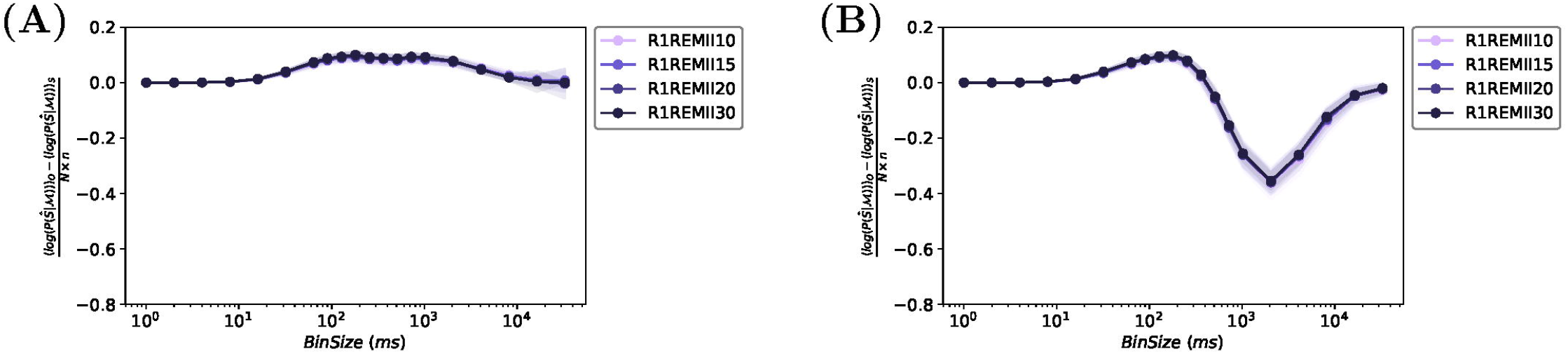
Normalized Log-Evidence per data point Vs. bin size when considering the difference of the number of the detected ICCs in original R1REMII dataset with the corresponding shuffled version. The plots are shown for two different methods of binning (A) based on the average firing rate and (B) based on the presence of spikes in bin. In both, the curves are normalized by the value of the *n*. This shows a universal behavior regardless of the cell’s population size that is considered in the analysis.

To this point, the results provide a good overview of how efficiently information is encoded by a minimal number of variables at around a certain value of bin size (∼10^2^), and the goodness of the inferred model around this value in comparison to other temporal resolutions.

### Orders of Interaction

According to the results of ICCs, one can expect that having components fewer than the number of cells signals us about the presence of multiple cells in each component. In order to visualize this better, we aim to understand how the number of cells in components/operators is affected by the temporal resolution that is used to analyze the data. To do so, we followed the procedure mentioned in section 0.6 for both components and operators inferred by MCMs procedure. The number of cells that are uniquely present in components/operators can be regarded as the order of interaction. For instance, if a component/operator is a product of three cells, then this component/operator is of the third order of interaction.

The difference heatmap plots for this matter show us temporal resolutions in which the orders of interaction are prominent in the dataset, and resolutions in which the bias introduces interactions of any order. Regardless of the binning method, results show that around 10^2^ milliseconds, we can observe the presence of the higher orders of interaction where there are components/operators with up to half of the population of the cells. However, this effect might be seen in large values of bin sizes, which, once again, might have been there due to lack of data points and false correlations among cells. This assumption can be supported by interactions of higher order introduced by bias around this bin size value (Ocher shades in Fig 4).

**Fig 4.**
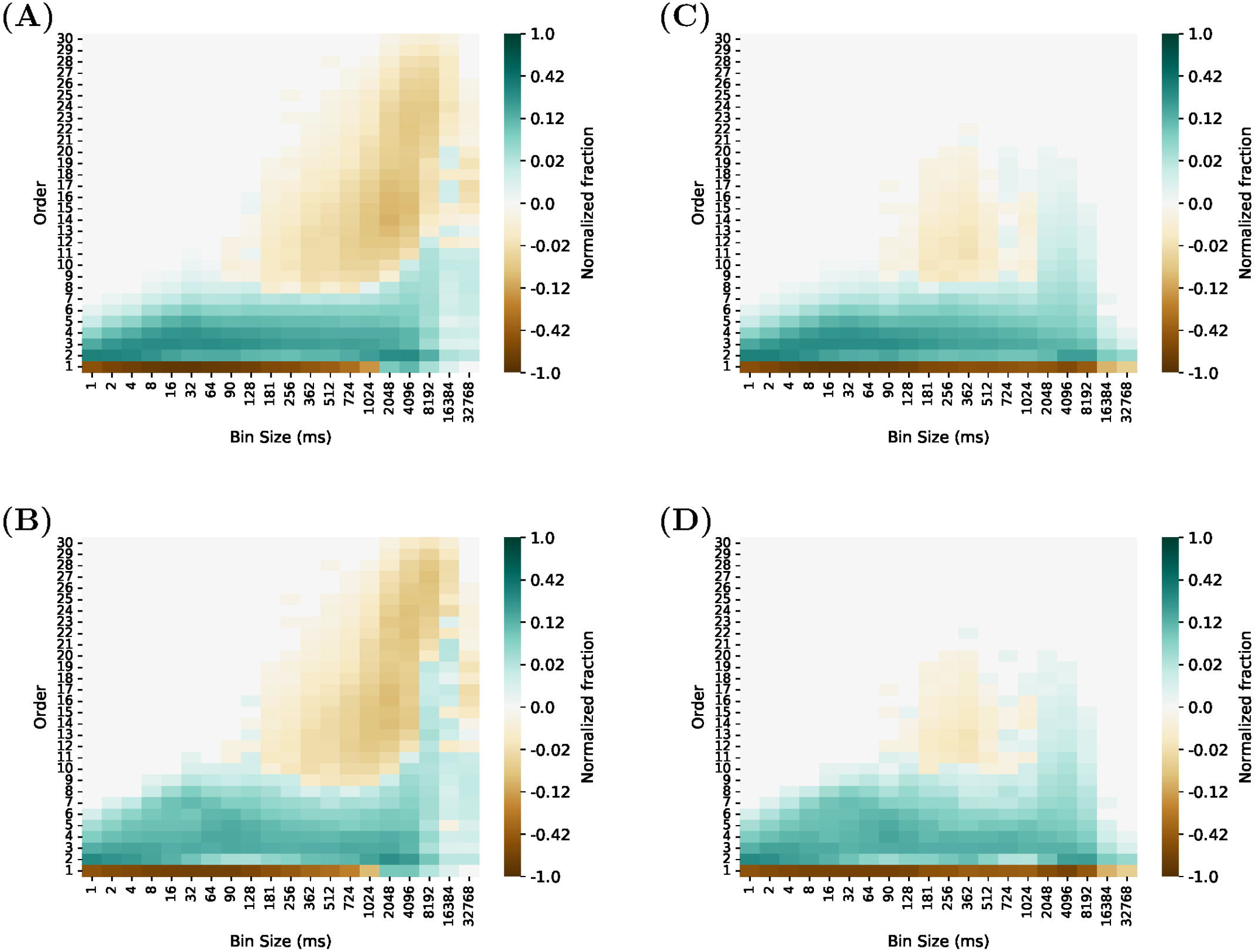
Normalized fraction of operators (panels A and C) / Components (panels B and D) that contain *n* number of cells (Order *n*) presented for Grid cell R1REMII data set. The plots are shown for two different methods of binning where the plots in the left column are based on the average firing rate and the ones in the right column are based on the presence of spikes in bin. Values are visualized in re-scaled format (explained in section 0.6) while preserving the original values on the color bar.

However, this results in the conclusion that the binning method based on presence of spikes is not as clear as the binning method based on average firing rate. One can confirm that when considering a great value of bin size, the decision of the activity based on presence of spikes leads to spike trains where almost all cells are active. This fact plus the lack of data points together lead to correlations which are not reliable.

### Distribution of Active Cells Across Different Bin Sizes

In order to have a general overview on the effect of temporal resolution on the activity of the cells, we attempt to understand the distribution of the active cells in different time bins. As mentioned in section 0.4, this analysis was done on original dataset, and we solely depend on the presence of cells regardless of the frequency of their activity. Results show interesting behaviors, one of which is the dominant fraction of bins at each bin size. This quantity follows a Sigmoid-like pattern which is dependent on the binning method; the Sigmoid-like pattern has saturation at different values of the fraction for active cells, in which by the saturation, we mean the value at the largest bin sizes considered in this paper. It is clear that if the bin size is equal to the size of the data points, both binning procedures result in the same value, which shows the activity of the cell by a single number. This observation confirms that the binning method based on the average firing rate leads to a sparser representation than the other method. In both methods, the sigmoid-like pattern shows almost the same behavior up to the point where 10% of the cells are active, which this fraction takes place around 10^2^ milliseconds and from then on, the dominant fraction of the bin size shows different values of active fractions. Furthermore, the general trend of the dominant fraction of bins overlaps or is not comparable, regardless of the dataset or the number of cells considered in the study (Fig 5).

**Fig 5.**
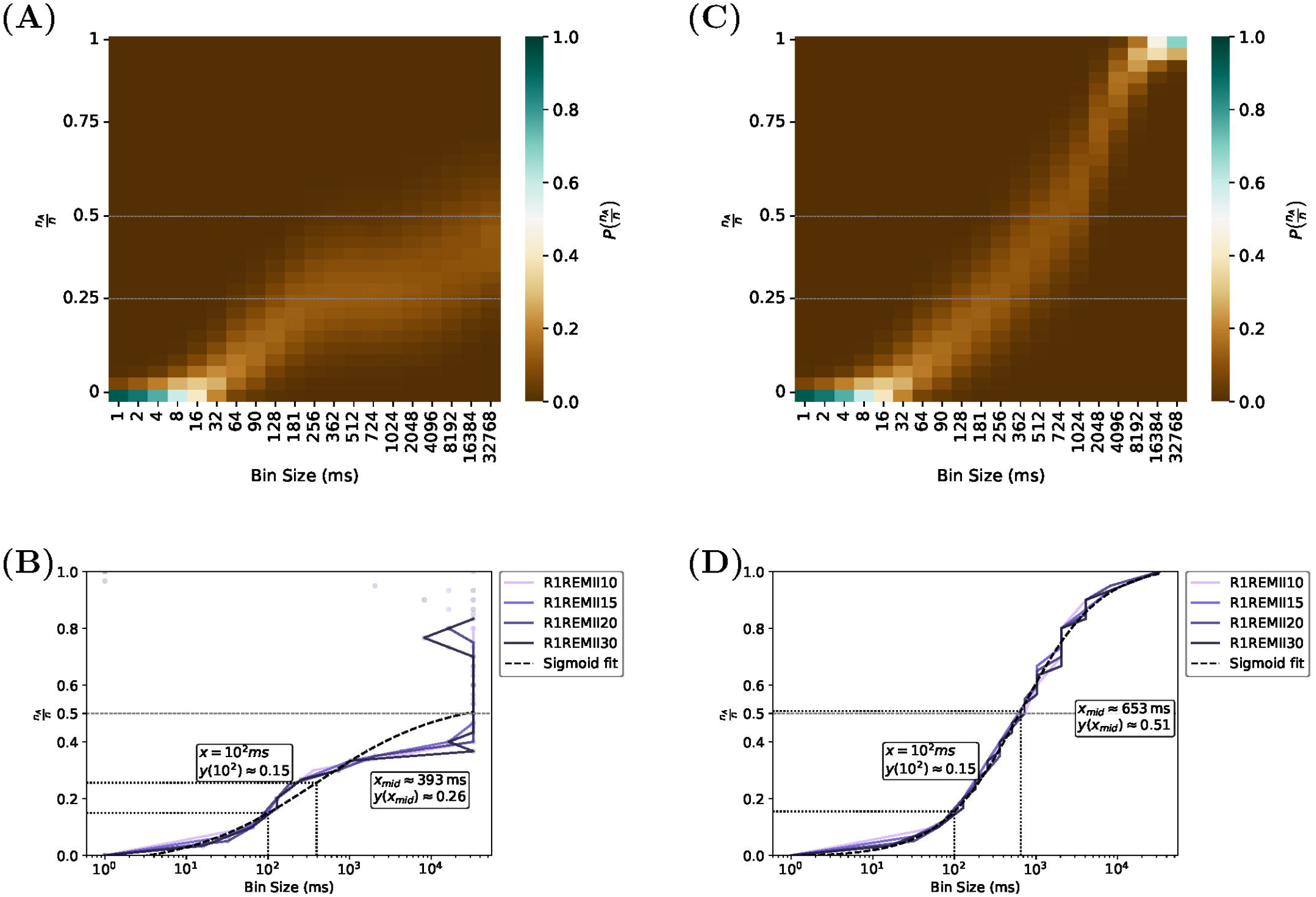
Fraction of bins with percentage of the active cells vs. bin size in R1REMII dataset with *n* = 30. The heat maps show the probability of the occurrence of each fraction of active cells after binning the data across different bin sizes, and the line plots show the most frequent/dominant fraction of active cells Vs. bin size for different populations of cells and Sigmoid fit on average of plots for different populations. The plots are shown for two different methods of binning where the plots in the left column are based on the average firing rate and the ones in the right column are based on the presence of spikes in bin. As can be seen, the most occurring fraction follows a Sigmoid-like pattern where the saturation of the most occurring fraction takes place around (A and B) 50% and (C and D) 100% of the total cells. The plot suggests where 50% of cells have the highest occurrence can be considered a key feature of the Sigmoid-like plots.

Furthermore, in both methods when the dominant numbers of the bins have 50% of the cells as active, it plays an important role. To be more precise, where the binning procedure is done based on the presence of a spike in bins, saturation takes place when almost 100% of cells are active, while when considering the binning method based on average firing rate, the saturation takes place at 50% of the fraction, which in both methods the saturation takes place around 10^4^ milliseconds. One can observe that 50% of the active fraction for the first case indicates the mid-point of the Sigmoid-like pattern (10^3^ milliseconds). This point corresponds to the point where the difference plot of log-evidence has minima. One can confirm the symmetrical distribution of the fraction of the active cells at this point, while smaller (larger) bin sizes lead to a bell-shaped distribution centered around (large) small number of cells, signaling the most frequent bins (Fig 6).

**Fig 6.**
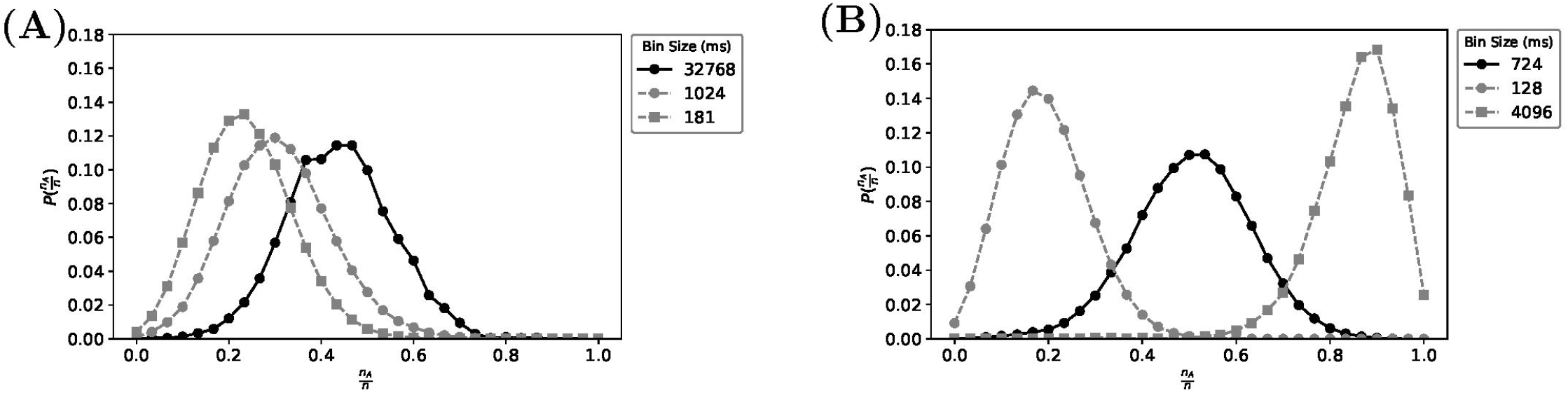
Distribution of the fraction of active cells Vs. bin size. Plots are shown for the binning method based on (A) firing rate (B) the presence of spikes in bin. This distribution is demonstrated for three bin sizes with a focus on the symmetrical pattern for the bin size where 50% of the bins have half of the cells active. One can observe that below this point, the distribution is centered around a small number of cells, while it is centered around a large number of cells for larger bin sizes in (B).

Clearly, based on the results, considering the temporal resolution in which the distribution of the active cells is centered around more than half of the cells (unsymmetrical distribution), leads to poor log-evidence and maximal introduction of the higher order interactions due to bias. On the other hand, before the bin size, where two methods of binning start to deviate when considering the distribution of the active cells (10^2^ milliseconds), is where the distributions are centered around almost 10% of the cells, showing rich behavior such as minimal number of components that encode information, maximal log-evidence and presence of the highest orders of interactions. Potentially this study can signal us about the most informative sparsity in neural networks too.

## Discussion

The presence of the higher order interactions and defining them in data recorded from complex interacting systems, has always been one of the central concerns. Traditional models attempt to infer the interactions by pre-defined models which limit the inference of potential orders of interaction which are absent in the model. By this attempt, the effort is to capture the most relevant information in a limited framework. However, we must admit that our knowledge of the underlying generative system is limited, for instance it is not clear exactly how the brain puts together the information from several neurons [35], and making any assumption about this system can potentially lead to false interpretations. On the other hand, it is not very clear whether or not prior assumptions capture all the present information in data. The story gets more complicated when scientists attempt to compare the importance of the higher order interactions across different co-variates. First and foremost, one must account for the present information in data which can give reliable information about the orders of interaction. One important underlying variable that determines the amount of information present in data, is the temporal resolution that the data is studied in. Clearly, at fine temporal resolutions, all the fine dynamical responses of the system’s elements are captured, while this variability is summarized into a single response at coarser temporal resolutions. Either of these hugely affects the correlation structure in data and consequently the information that one can obtain from the data [36]. This important factor has been studied extensively when it comes to extracting information from the recording of one system’s element [37–46]. However, it is understood that the activity of every individual cell is not isolated from other cells and the stimulus is not the only factor in how the neuron responds [47, 48]. When it comes to the brain, every individual cell receives around 10000 synaptic connections from other neurons [49, 50]. Therefore, it is very important to address the effect of the temporal resolution when it comes to the study of a population of elements.

This paper demonstrates the key role of the temporal resolution in encoding of the information and the importance of the higher orders of interaction. Based on the inference method used in this study, one can systematically infer all the present interactions solely based on the data without any prior assumption on interactions of the underlying model. This inference gives us a good overview of the number of components that are needed in order to encode the information efficiently and orders of interactions. These two factors are highly affected by the temporal resolution at which the data is studied. This study showcased two temporal resolutions as the most important ones. One takes place at fine resolution (∼10^2^ ms) and another one in coarse resolution. The former point is associated with a minimal number of independent components, maximal un/normalized log evidence per data point, rich correlational structure (up to the 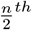 order of interaction) and sparse representation of the network where the highest fraction of the bins include ∼10% active cells. The latter point is associated with distributed representations where the distribution of the active cells for each bin is drawn from a normal distribution centered around 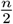. This is a point where false correlations appear to be most dominant due to the bias of limited data (yellow region in Fig 8 and Fig 9), the minimal un/normalized log evidence per data point, and a minimal number of the independent components when encoding shuffled data.

Although distributed representations show robustness and high capacity in storing and retrieving information [51], it comes with redundancy when dealing with inference in limited data, which is confirmed in this study. This contrast can be observed where higher orders of interaction are present (Fig 4) despite the poor inferred models (Fig 3) (note that the minimas of the log evidence plots happen at the time resolutions where distributed representation occurs). Another important conclusion is that the comparison of the orders of interaction should be made according to the temporal resolution that the data is studied. For instance, one cannot simply claim that “second order of interaction is always prominent”, rather one can state that “second order interaction is prominent when the data is considered at the finest scale”. For our case study, one can observe that for instance, the second order of interaction is more prominent than 6^*th*^ order of interaction when the data is studied at 1*ms*. However, around 100*ms* this prominence is reversed. This result can be confirmed in the second and sixth rows of the heatmaps Fig 4 at the specified temporal resolution or bin size in milliseconds. Therefore, when comparing the importance of orders of interaction regarding internal or external co-variates, one must first and foremost account for temporal resolution (Check S1 Text).

Furthermore, this study might shed light on the optimal time resolution that one might study the data in order to obtain the maximal compressed information. As per Shannon’s idea, the compression of the data is best done when the data is presented with a minimal number of variables [52, 53]. This study shows that for our case study data, there exists a temporal resolution in which informational compressibility is maximal.

Our study can be used when studying complex interacting systems in an efficient manner, particularly when limited data is available. Based on this framework, one can first determine the most and least informative temporal resolutions and continue with further analysis in an accurate and systematic setting.

## Conclusion

This study shed light on the importance of the temporal resolution when it comes to studying complex interacting systems based on limited dataset. By employing the framework of the Minimally Complex Models (MCMs), we systematically inferred the structure of interactions in complex interacting systems with no prior assumption of the structure across different time resolutions. Considering different methods of binning, the results show the importance of two pivotal temporal resolutions; one around 100 millisecond and the other where the network has distributed representation. The former pivot point is characterized by a minimal number of independent complete components, maximal un/normalized log evidence per data points, dominant presence of the higher orders of interaction and sparse representation of the network (∼10%). While the latter pivot point is characterized by minimal number of independent complete components for shuffled data, minimal un/normalized log evidence per data points, dominant presence of the higher orders of interaction in shuffled data and distributed representation of the network (∼50%). Normalized log evidence per data points as a parameter which is independent of the system size, can be used as a universal ruler for determining these pivot points. This study emphasizes that the amount of information in data is highly affected by the temporal resolution and consequently the relevance of the *n*^*th*^ order of interaction is dependent on the temporal resolution. Our findings underscore the importance of accounting for the temporal resolution when it comes to accurate interpretation of data from interacting systems, especially when the attempt is comparing different states of co-variates. In conclusion, this work provides a systematic approach in order to understand the underlying pattern of generative model in complex interacting systems based on limited data, with implications in different fields such as neuroscience and beyond.

## Supporting information

### Temporal resolution affects comparison of the significance of the orders of interactions

In this part, the dependency of the higher orders of interaction on external and internal co-variates is addressed. Based on the results, there is no absolute statement that one can make about the prominence of orders of interaction in different experimental states or modules. This prominence is highly dependent on the temporal resolution that each order of interaction is studied. Having the temporal resolution as a fixed parameter, one can attempt to compare different co-variates. For instance, for rat R on day II, considering temporal resolutions ∈ {1, ∼ 10^2^}milliseconds for both binning methods, we can see patterns that are unique to these resolutions. If we namely associate the difference of the order’s probability (*P* (*Order*)_*O*_ ™ *P* (*Order*)_*S*_) as significance, when considering 1 millisecond temporal resolution, REM shows lower significance for single cell operators than OF and SWS, while for operators with three cells, REM has the highest significance (S1 Fig). On the other hand, on the scale of 128 millisecond, operators with first and second orders show the highest significance for REM, followed by slightly more significance of the OF than SWS, while this pattern is reversed for operators with order greater than 2, i.e. SWS *>* OF *>* REM (S2 Fig). These results are consistent in both binning methods. When considering the component’s order, one can more or less obtain the same conclusions, with the difference that the reverse of the patterns takes place in higher orders of interaction in comparison with operators (S3 Fig and S4 Fig). This can potentially arise from the richer statistics of operators. In general, results confirm the important role of the temporal resolution on the significance of the co-variates for different orders of interaction.

**S1 Fig.**
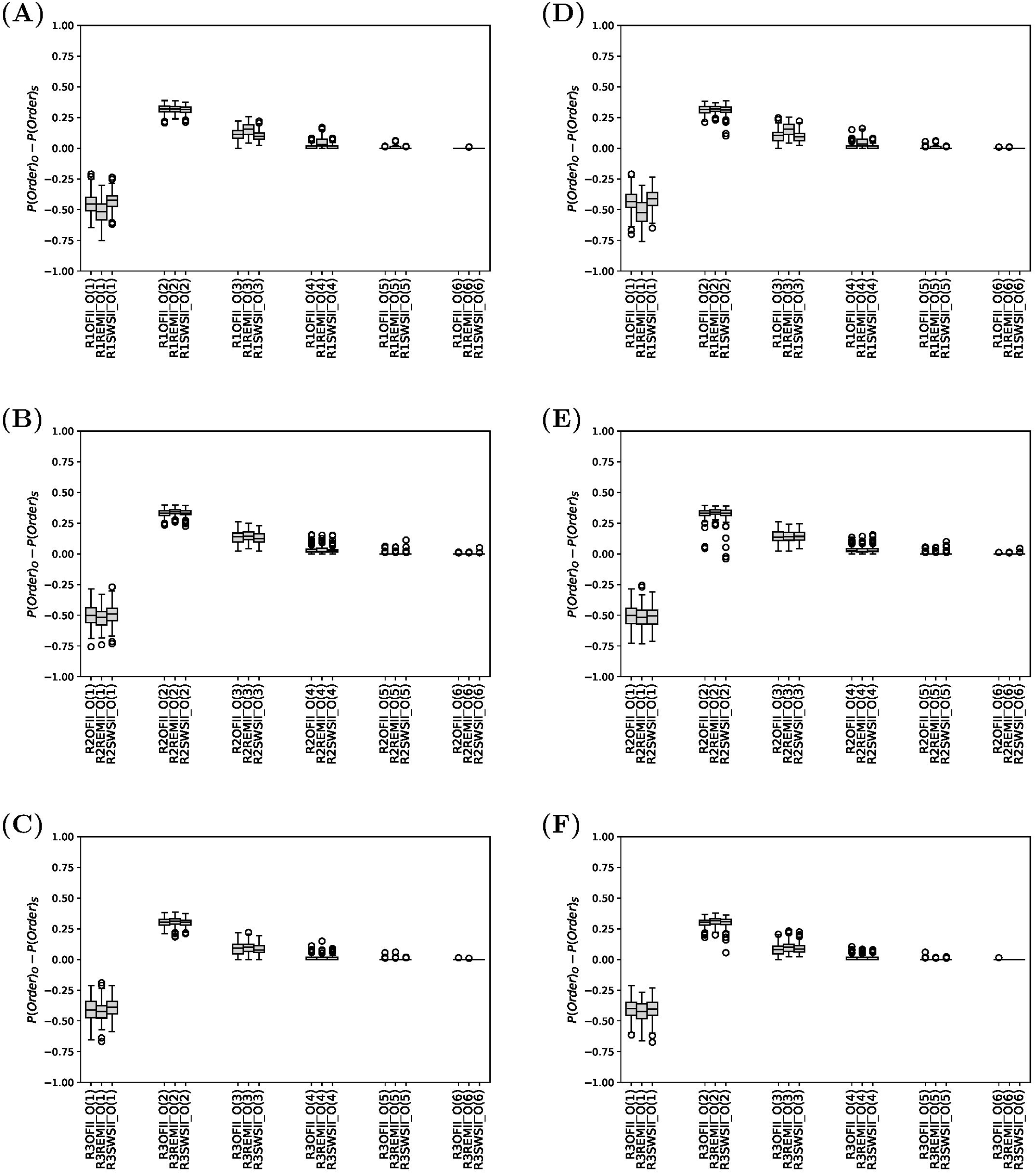
Comparison of the different experimental co-variates for each module of rat R on day II considering orders of interaction in operators (1 millisecond). The left (right) column belongs to the binning method based on average firing rate (presence of spikes).

**S2. Fig.**
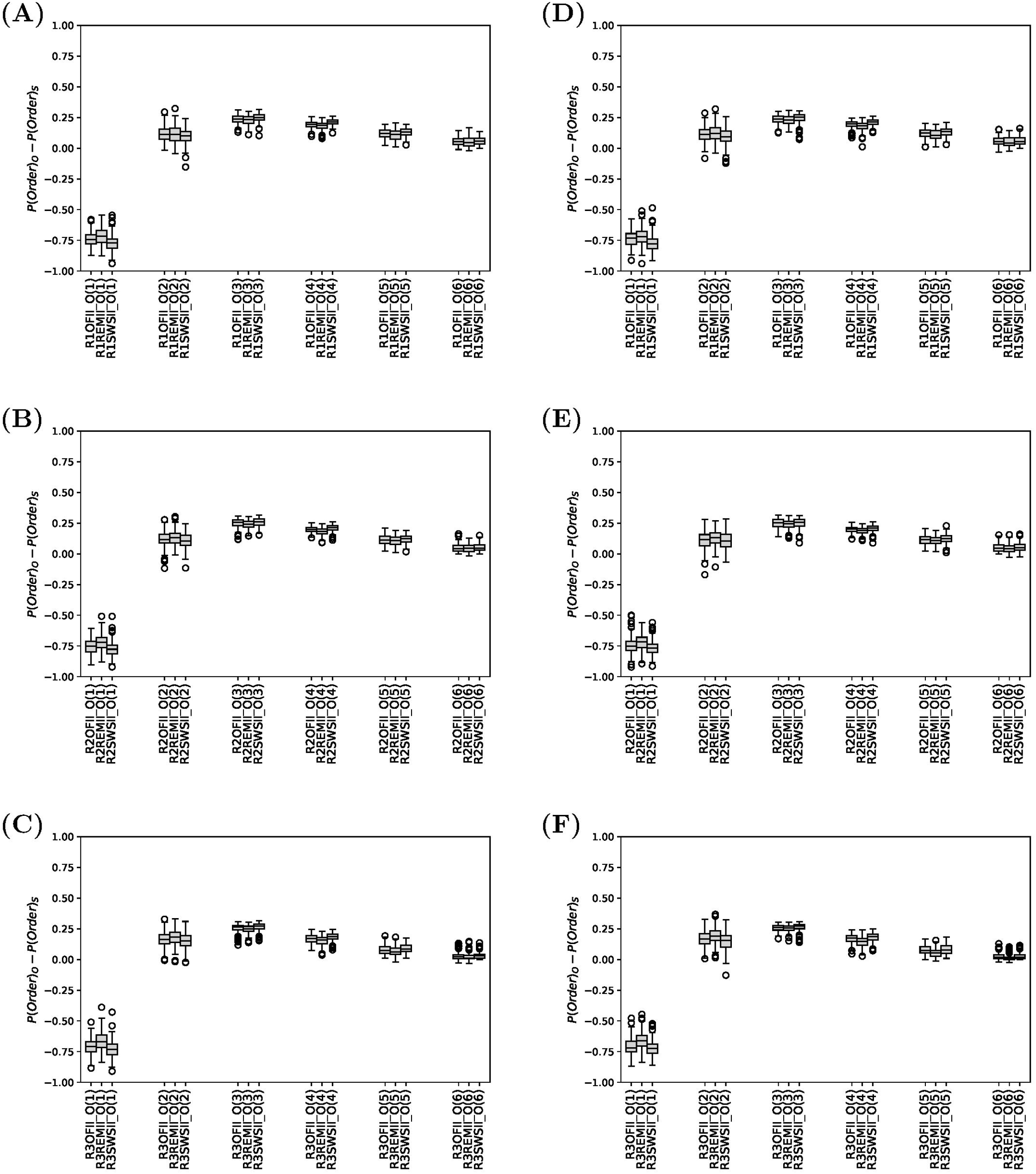
Comparison of the different experimental co-variates for each module of rat R on day II considering orders of interaction in operators (128 millisecond). The left (right) column belongs to the binning method based on average firing rate (presence of spikes).

**S3. Fig.**
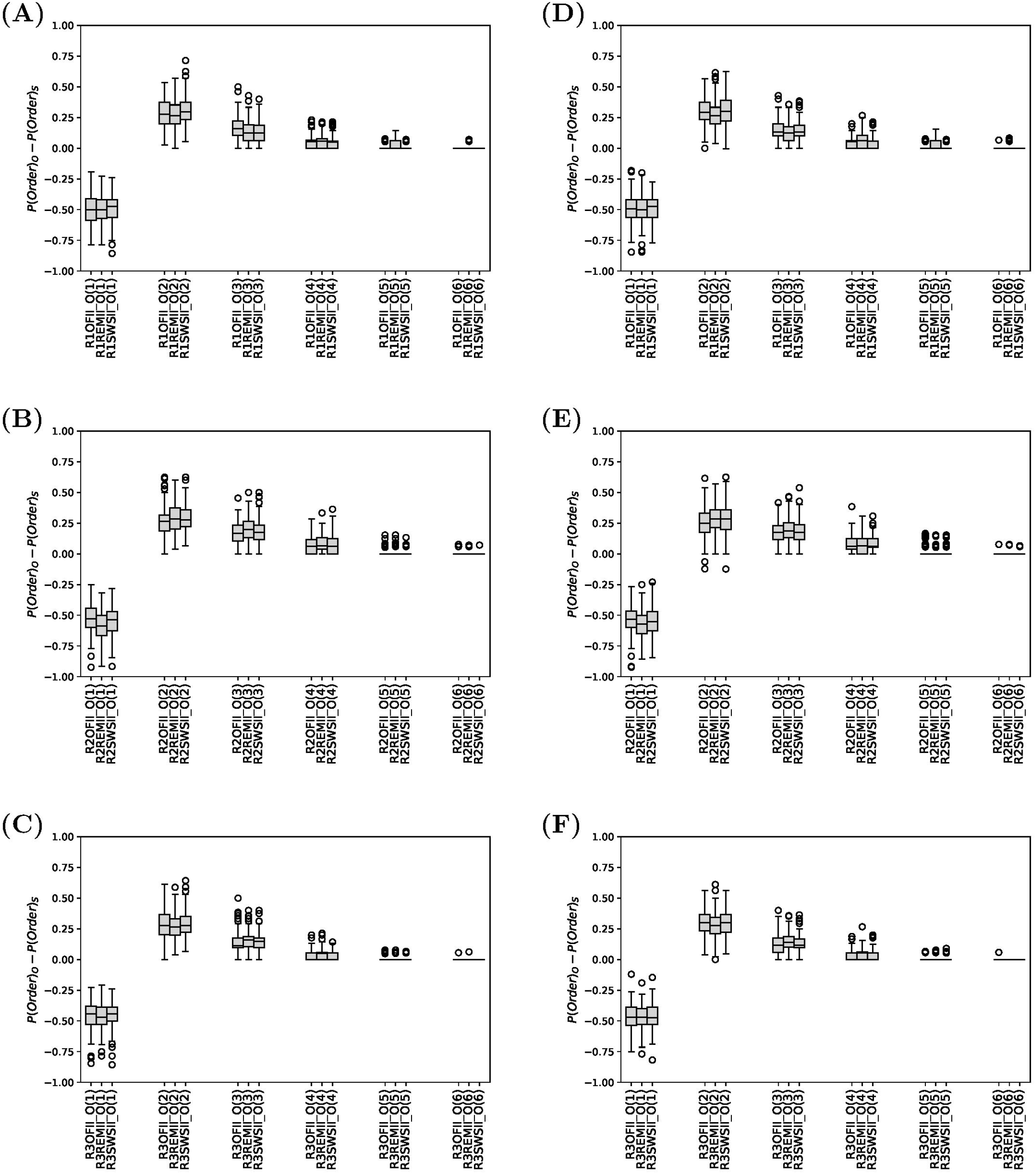
Comparison of the different experimental co-variates for each module of rat R on day II considering orders of interaction in components (1 millisecond). The left (right) column belongs to the binning method based on average firing rate (presence of spikes).

**S4. Fig.**
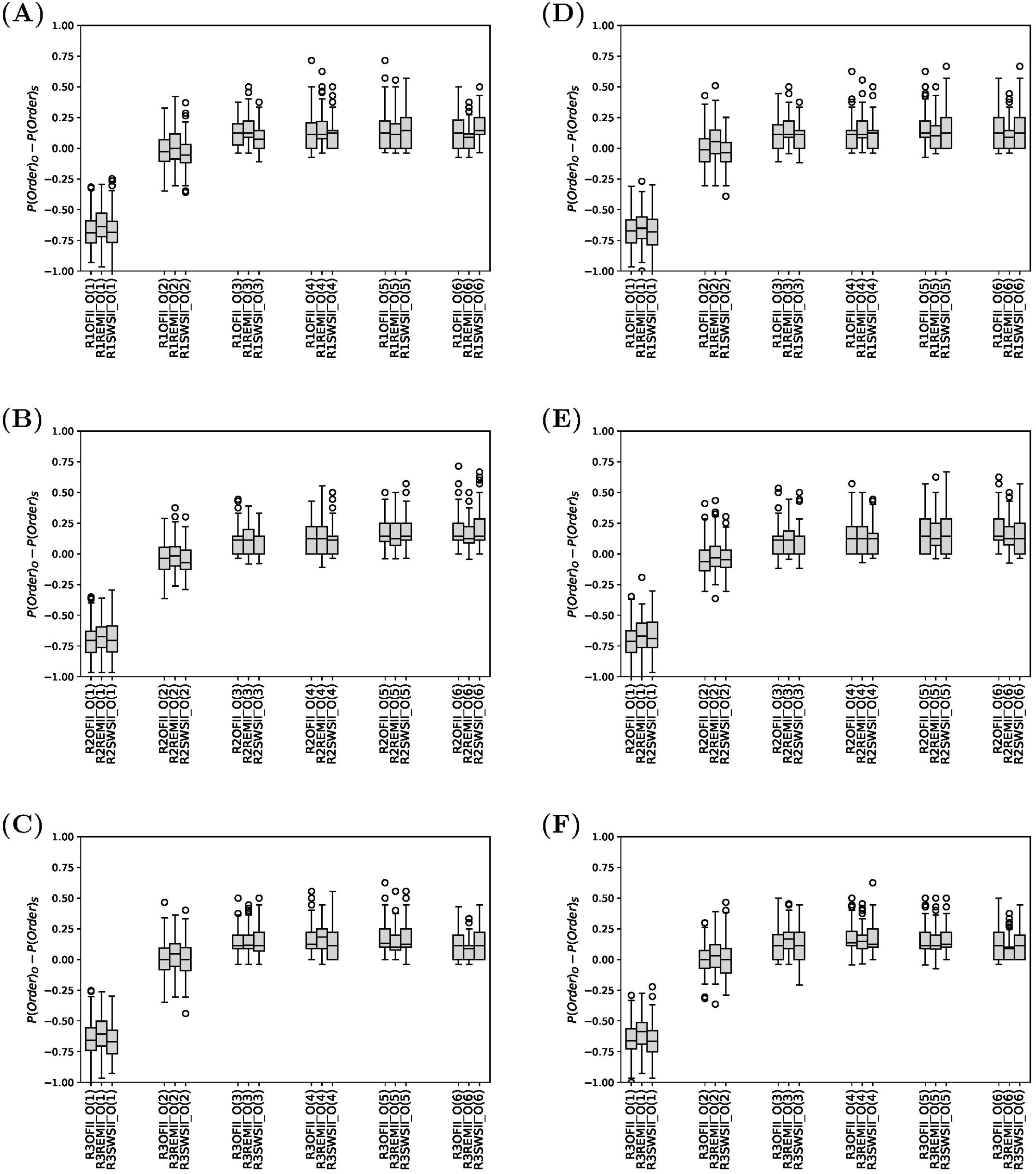
Comparison of the different experimental co-variates for each module of rat R on day II considering orders of interaction in components (128 millisecond). The left (right) column belongs to the binning method based on average firing rate (presence of presence of spikes).

## Acknowledgments

The author would like to thank Yasser Roudi and Matteo Marsili for valuable discussions and insights regarding the methodology, analysis methods, and results. The author also thanks Clélia de Mulatier for valuable discussions, which led to clarification on minimally complex models.

## Data Availability Statement

All data, codes, and results are openly available in the Open Science Framework (OSF) repository at: https://osf.io/jn6m5/?view_only=33e6056f2f6a4ab396504344ccf9b57a.

The original dataset and data extraction code are from Gardner et al., Nature (2022), and can be accessed at: https://github.com/erikher/GridCellTorus and https://figshare.com/articles/dataset/Toroidal_topology_of_population_activity_in_grid_cells/16764508.

## References

1. Yu S, Yang H, Nakahara H, Santos GS, Nikolić D, Plenz D. Higher-order interactions characterized in cortical activity. Journal of neuroscience. 2011;31(48):17514–17526.

2. Ince RA, Montani F, Arabzadeh E, Diamond ME, Panzeri S. On the presence of high-order interactions among somatosensory neurons and their effect on information transmission. In: Journal of Physics: Conference Series. vol. 197. IOP Publishing; 2009. p. 012013.

3. Montani F, Ince RA, Senatore R, Arabzadeh E, Diamond ME, Panzeri S. The impact of high-order interactions on the rate of synchronous discharge and information transmission in somatosensory cortex. Philosophical Transactions of the Royal Society A: Mathematical, Physical and Engineering Sciences. 2009;367(1901):3297–3310.

4. Martignon L, Deco G, Laskey K, Diamond M, Freiwald W, Vaadia E. Neural coding: higher-order temporal patterns in the neurostatistics of cell assemblies. Neural computation. 2000;12(11):2621–2653.

5. Ganmor E, Segev R, Schneidman E. Sparse low-order interaction network underlies a highly correlated and learnable neural population code. Proceedings of the National Academy of sciences. 2011;108(23):9679–9684.

6. Zylberberg J, Shea-Brown E. Input nonlinearities can shape beyond-pairwise correlations and improve information transmission by neural populations. Physical Review E. 2015;92(6):062707.

7. Schneidman E, Berry MJ, Segev R, Bialek W. Weak pairwise correlations imply strongly correlated network states in a neural population. Nature. 2006;440(7087):1007–1012.

8. Kuhn A, Aertsen A, Rotter S. Higher-order statistics of input ensembles and the response of simple model neurons. Neural computation. 2003;15(1):67–101.

9. Amari Si, Nakahara H, Wu S, Sakai Y. Synchronous firing and higher-order interactions in neuron pool. Neural computation. 2003;15(1):127–142.

10. Cunningham JP, Yu BM. Dimensionality reduction for large-scale neural recordings. Nature neuroscience. 2014;17(11):1500–1509.

11. Panzeri S, Harvey CD, Piasini E, Latham PE, Fellin T. Cracking the neural code for sensory perception by combining statistics, intervention, and behavior. Neuron. 2017;93(3):491–507.

12. Cubero RJ, Marsili M, Roudi Y. Multiscale relevance and informative encoding in neuronal spike trains. Journal of computational neuroscience. 2020;48:85–102.

13. Cubero RJ, Jo J, Marsili M, Roudi Y, Song J. Statistical criticality arises in most informative representations. Journal of Statistical Mechanics: Theory and Experiment. 2019;2019(6):063402. doi:10.1088/1742-5468/ab16c8.

14. Panzeri S, Schultz SR, Treves A, Rolls ET. Correlations and the encoding of information in the nervous system. Proceedings of the Royal Society of London Series B: Biological Sciences. 1999;266(1423):1001–1012.

15. Lambiotte R, Rosvall M, Scholtes I. From networks to optimal higher-order models of complex systems. Nature physics. 2019;15(4):313–320.

16. Strong SP, Koberle R, Van Steveninck RRDR, Bialek W. Entropy and information in neural spike trains. Physical review letters. 1998;80(1):197.

17. Owen LL, Chang TH, Manning JR. High-level cognition during story listening is reflected in high-order dynamic correlations in neural activity patterns. Nature Communications. 2021;12(1):5728.

18. Parastesh F, Mehrabbeik M, Rajagopal K, Jafari S, Perc M. Synchronization in Hindmarsh–Rose neurons subject to higher-order interactions. Chaos: An Interdisciplinary Journal of Nonlinear Science. 2022;32(1).

19. Trettel SG, Trimper JB, Hwaun E, Fiete IR, Colgin LL. Grid cell co-activity patterns during sleep reflect spatial overlap of grid fields during active behaviors. Nature neuroscience. 2019;22(4):609–617.

20. Gardner RJ, Lu L, Wernle T, Moser MB, Moser EI. Correlation structure of grid cells is preserved during sleep. Nature neuroscience. 2019;22(4):598–608.

21. Li Q, Yu S, Madsen KH, Calhoun VD, Iraji A. Higher-order organization in the human brain from matrix-based rényi’s entropy. In: 2023 IEEE International Conference on Acoustics, Speech, and Signal Processing Workshops (ICASSPW). IEEE; 2023. p. 1–5.

22. Valenti S, Sparacino L, Pernice R, Marinazzo D, Almgren H, Comelli A, et al. Assessing High-Order Interdependencies Through Static O-Information Measures Computed on Resting State fMRI Intrinsic Component Networks. In: International Conference on Image Analysis and Processing. Springer; 2022. p. 386–397.

23. Latham PE, Nirenberg S. Synergy, redundancy, and independence in population codes, revisited. Journal of Neuroscience. 2005;25(21):5195–5206.

24. Reynolds JH, Chelazzi L. Attentional modulation of visual processing. Annu Rev Neurosci. 2004;27(1):611–647.

25. Shlens J, Field GD, Gauthier JL, Grivich MI, Petrusca D, Sher A, et al. The structure of multi-neuron firing patterns in primate retina. Journal of Neuroscience. 2006;26(32):8254–8266.

26. Montani F, Phoka E, Portesi M, Schultz SR. Statistical modelling of higher-order correlations in pools of neural activity. Physica A: Statistical Mechanics and its Applications. 2013;392(14):3066–3086.

27. Bohté SM, Spekreijse H, Roelfsema PR. The effects of pair-wise and higher-order correlations on the firing rate of a postsynaptic neuron. Neural Computation. 2000;12(1):153–179.

28. Nakahara H, Amari Si. Information-geometric measure for neural spikes. Neural computation. 2002;14(10):2269–2316.

29. Olsen VK, Whitlock JR, Roudi Y. The quality and complexity of pairwise maximum entropy models for large cortical populations. PLOS Computational Biology. 2024;20(5):e1012074.

30. Pillow JW, Shlens J, Paninski L, Sher A, Litke AM, Chichilnisky E, et al. Spatio-temporal correlations and visual signalling in a complete neuronal population. Nature. 2008;454(7207):995–999.

31. Benucci A, Verschure PF, König P. Existence of high-order correlations in cortical activity. Physical Review E. 2003;68(4):041905.

32. Roudi Y, Nirenberg S, Latham PE. Pairwise maximum entropy models for studying large biological systems: when they can work and when they can’t. PLoS computational biology. 2009;5(5):e1000380.

33. de Mulatier C, Mazza PP, Marsili M. Statistical inference of minimally complex models. arXiv preprint arXiv:200800520. 2020;.

34. Gardner RJ, Hermansen E, Pachitariu M, Burak Y, Baas NA, Dunn BA, et al. Toroidal topology of population activity in grid cells. Nature. 2022;602(7895):123–128.

35. Ince RA, Senatore R, Arabzadeh E, Montani F, Diamond ME, Panzeri S. Information-theoretic methods for studying population codes. Neural Networks. 2010;23(6):713–727.

36. Von der Malsburg C. The what and why of binding: the modeler’s perspective. Neuron. 1999;24(1):95–104.

37. Mahalle PN, Ambritta P N, Sakhare SR, Kulkarni AP. Data Science Problems. In: Foundations of Mathematical Modelling for Engineering Problem Solving. Springer; 2023. p. 87–141.

38. Irie MS, Spin-Neto R, Borges JS, Wenzel A, Soares PBF. Effect of data binning and frame averaging for micro-CT image acquisition on the morphometric outcome of bone repair assessment. Scientific Reports. 2022;12(1):1424.

39. Scott DW. On optimal and data-based histograms. Biometrika. 1979;66(3):605–610.

40. Shimazaki H, Shinomoto S. A recipe for optimizing a time-histogram. Advances in neural information processing systems. 2006;19.

41. Shimazaki H, Shinomoto S. A method for selecting the bin size of a time histogram. Neural computation. 2007;19(6):1503–1527.

42. Shimazaki H, Shinomoto S. Kernel bandwidth optimization in spike rate estimation. Journal of computational neuroscience. 2010;29:171–182.

43. Freedman D, Diaconis P. On the histogram as a density estimator: L 2 theory. Zeitschrift für Wahrscheinlichkeitstheorie und verwandte Gebiete. 1981;57(4):453–476.

44. Wand M. Data-based choice of histogram bin width. The American Statistician. 1997;51(1):59–64.

45. Omi T, Shinomoto S. Optimizing time histograms for non-Poissonian spike trains. Neural computation. 2011;23(12):3125–3144.

46. Heidarieh S, Jahed M, Ghazizadeh A. Variable bin size selection for periestimulus time histograms (PSTH) with minimum mean square error criteria. BMC Neuroscience. 2015;16:1–2.

47. Abeles M, Bergman H, Margalit E, Vaadia E. Spatiotemporal firing patterns in the frontal cortex of behaving monkeys. Journal of neurophysiology. 1993;70(4):1629–1638.

48. Li CL. Synchronization of unit activity in the cerebral cortex. Science. 1959;129(3351):783–784.

49. Paisley A, Summerlee A. Relationships between behavioural states and activity of the cerebral cortex. Progress in neurobiology. 1984;22(2):155–184.

50. Legendy C, Salcman M. Bursts and recurrences of bursts in the spike trains of spontaneously active striate cortex neurons. Journal of neurophysiology. 1985;53(4):926–939.

51. Rolls E, Treves A. Neural networks and brain function. Oxford university press; 1997.

52. Shannon CE. A mathematical theory of communication. The Bell system technical journal. 1948;27(3):379–423.

53. Shannon CE. Communication in the presence of noise. Proceedings of the IRE. 1949;37(1):10–21.

